# A Rapid and Highly Sensitive CRISPR Assay Utilizing Cas12a Orthologs for the Detection of Novel Duck Reovirus

**DOI:** 10.1101/2025.03.27.645718

**Authors:** Yuan Wang, Lanting Fu, Sheng Li, Dagang Tao, Ping Gong, Yu Yang, Jinxue Ruan, Shengsong Xie, Cui Wang, Daqian He

## Abstract

The novel duck reovirus (NDRV) disease presents a significant threat to the poultry industry due to the absence of effective therapeutic measures. As a result, there is an urgent need to develop innovative rapid diagnostic methods for early virus detection. In this study, we developed a Rapid Visual CRISPR Assay to detect the NDRV S3 gene using novel Cas12a orthologs. Specifically, we compared the performance of two candidates, Gs12-16 and Gs12-18, in detecting the NDRV S3 gene to identify a highly sensitive and efficient CRISPR-based diagnostic method. Our results demonstrated that both Gs12-16 and Gs12-18 exhibited strong *cis*- and *trans*-cleavage activities for classical “TTTV” protospacer adjacent motif (PAM)-containing targets *in vitro*, although they required different reaction temperatures. Notably, Gs12-18 showed relatively higher activity for dsDNA targets compared to Gs12-16, indicating that Gs12-18 is more suitable for CRISPR-based nucleic acid detection applications. To leverage these properties, we integrated Gs12-18 with loop-mediated isothermal amplification (LAMP) technology to establish a LAMP-CRISPR/Gs12-18-mediated method for detecting the NDRV S3 gene. This approach enables highly sensitive and visually detectable on-site identification of the NDRV S3 gene, achieving a sensitivity of 38 copies per reaction. Our LAMP-CRISPR/Gs12-18-based method can be utilized for highly sensitive detection of NDRV nucleic acids.

## 1 Introduction

Novel duck reovirus disease (NDR), caused by a novel duck reovirus (NDRV), poses a significant threat to the global duck farming industry^[1]^. First identified in southern China in 2005, this highly pathogenic virus induces severe immunosuppressive symptoms characterized by hemorrhagic necrosis in immune organs such as the liver and spleen^[2,3]^. The nationwide outbreak in 2011 further highlighted its impact, affecting a broader range of duck species with increased severity compared to the previously reported Muscovy duck reovirus (MDRV) ^[4,5]^.

Current detection methods for NDRV include the agar gel precipitation test (AGPT), Enzyme linked immunosorbent assay (ELISA)^[6,7]^, RT-PCR^[8]^, RT-qPCR^[9,10]^, Reverse Transcription-Recombinase Polymerase Amplification (RT-RPA)^[11,12]^, and RT-loop mediated isothermal amplification (RT-LAMP)^[13]^. While AGPT is simple and rapid, it suffers from low sensitivity and cannot differentiate between multiple serotypes^[14]^. ELISA offers higher sensitivity and specificity but faces challenges in early virus detection due to limited antibody presence during the initial infection stages^[6,7]^. Nucleic acid detection techniques like PCR-based methods require specialized equipment and trained personnel, and some exhibit low specificity and sensitivity, making field-based rapid detection difficult^[8-10]^. Although isothermal amplification techniques like RT-RPA and RT-LAMP provide high sensitivity, they are prone to non-specific amplification resulting in false positive results^[11-13]^. Therefore, there is an urgent need to develop a rapid, highly sensitive, and easy-to-perform assay for the field/point-of-care detection of novel duck reoviruses.

In recent years, the CRISPR/Cas system combined with isothermal amplification reaction-mediated nucleic acid detection technology has shown great potential^[15,16,17,18]^. This method offers high sensitivity and specificity, enabling rapid and accurate on-site detection. The principle involves CRISPR/Cas12 protein, guided by CRISPR RNA (crRNA), recognizing and cleaving specific target DNA, activating its *trans*-cleavage activity to efficiently degrade non-specific single-stranded DNA (ssDNA) within the system^[19]^. Our research team previously developed a series of Cas12a-based nucleic acid detection platforms named RAVI-CRISPR[20] or multiplex RAVI-CRISPR^[21]^. The first generation was designed by optimizing single-stranded DNA-fluorophore-quencher (ssDNA-FQ) technology, enabling visual detection without specialized instruments^[20,22]^. The second generation introduced a dual-gene diagnostic system leveraging the distinct *trans*-cleavage activities of Cas12a and Cas13a enzymes^[21]^.

In this study, we conducted a systematic comparison and evaluation of the activity characteristics of two Cas12a orthologs, Gs12-16 and Gs12-18 enzymes identified in our previous study, to establish a RAVI-CRISPR-based nucleic acid detection assay mediated by these novel Cas12a homologs^[23]^. Our results demonstrated that both enzymes exhibited *cis*- and *trans*-cleavage activities, with optimal temperature ranges of 37–45 °C and 25–60 °C, respectively. Both exhibited cleavage activity within 5 minutes, with Gs12-18 showing a broader temperature range and a more robust *trans*-cleavage effect. Based on these findings, we developed a new detection technique for NDRV using the LAMP-CRISPR/Gs12-18 system, achieving high sensitivity and specificity for viral nucleic acid detection. This method facilitates early prevention and control of NDRV, offering a promising solution for the global duck farming industry. Future work will further optimize the RAVI-CRISPR platform and explore additional CRISPR tools to enhance its versatility and effectiveness in nucleic acid detection.

## 2 Materials and Methods

### 2.1 Sequence conservation analysis of two candidate Cas12a orthologs

We evaluated the evolutionary relationship in two Cas12a homologs, namely Gs12-16 and Gs12-18, identified in our previous study ^[23]^. PSI-BLAST was used to determine the Gs12-16 and Gs12-18 protein sequences from an existing dataset of five known Cas12a proteins: LbCas12a, FnCas12a, AsCas12a, MbCas12a and PiCas12a. The resulting seven Cas12a protein sequences were subjected to multiple sequence alignments using Geneious Prime (www.geneious.com/features/prime), resulting in a comprehensive sequence conservation analysis of these orthologs.

### 2.2 Purification of Gs12-16 and Gs12-18 proteins

We obtained purified Gs12-16 and Gs12-18 proteins by first synthesizing their DNA sequences (Table S4), cloning them into the pET28a expression vector, and transforming the plasmids into chemically competent *E. coli* BL21 (λDE3) (Cat#C2527H, NEB) according to the manufacturer’s protocol. The transformed *E. coli* was plated onto <u>Kanamycin</u> agar plates supplemented with 50 µg/mL and incubated overnight at 37 °C. Single colonies from the overnight cultures were resuspended in 300 mL of broth media with antibiotic and incubated at 37 °C with shaking at 200 rpm until an optical density (OD_600_) of 0.6 was reached. Protein expression was induced by adding 0.5 mM Isopropyl β-D-Thiogalactoside (IPTG) into the broth culture and incubating at 18 °C for 16 hours.

The bacterial precipitates were collected and resuspended in a lysate containing 300 mM NaCl, 2 mM MgCl_2_, 5 mM HEPES, 20 mM imidazole, and 1 mM DTT. Subsequent sonication and centrifugation at 8,000 rpm for 30 minutes at 4 °C yielded a supernatant that was incubated with Nickel Affinity Chromatography Column Resin (Huiyan Bio, HQ060312025M) at 4 °C for 2 hours. The sample was then passed through the gravity column three times, and the supernatant was discarded. A 10-fold volume of wash solution (500 mM NaCl, 10 mM tris, and 30 mM imidazole) was added, followed by elution of the target proteins with an elution solution (500 mM NaCl, 10 mM tris, and 300 mM imidazole). The collected proteins were verified by SDS-PAGE, and those with minimal heteroproteins were concentrated using a 30 kDa concentration column and preserved in a storage solution (500 mM NaCl, 20 mM sodium acetate, 0.1 mM EDTA, and 50% glycerol) at -80 °C for future use. Protein quantification was performed using the BCA protein assay kit (Cat#P0012, beyotime).

### 2.3 Nucleic acid preparation

Sequences from 26 different genotypes of NDRV strains were retrieved from GenBank (accession numbers: MK955825, MH510263, MH510253, KT861595, ON009445, OM930743, MW924630, KJ879932, MT829206, KF163099, KF163098, KF163097, KF163096, KF163095, JQ866923, JX478268, MW435676, MW435673, MW435672, MW435671, MW435668, MW435667, JX826588, MN747012, HM591301, and GQ888710). We used DNAMAN software to compare multiple sequences and analyze the S3 genes. A partial segment of the S3 gene from the DRVGX-Y7 strain was synthesized by Tsingke and designated as NDRV S3 (Table S3). The synthesized NDRV S3 plasmid served as a template for PCR amplification using primers NDRV-F (5’-TGTTGCACTCAGTGCTGTGG-3’) and NDRV-R (5’-AATTGACCAGTGATGCCAAC-3’). This yielded the double-stranded DNA (dsDNA) of the NDRV S3 gene, which was used in subsequent studies.

### 2.4 Preparation of crRNAs

Using the NDRV S3 gene as the target sequence, we designed 16 crRNAs for classical protospacer adjacent motif (PAM, such as TTTV or TTV, where V represents A, C, or G) targets at distinct sites, and 12 crRNAs for non-classical PAM targets at different locations (Table S1). We amplified and purified the crRNA *in vitro* transcript templates using the T7-F forward and reverse primers. The resulting PCR product was transcribed overnight at 37 °C using the HiScribe T7 High Yield RNA Synthesis Kit (NEB, Cat. No. E2040S). The synthesized RNA was subsequently purified using the Monarch RNA Cleanup Kit (NEB, Cat. No. T2040S), and its concentration was determined using a NanoDrop™ 2000 Spectrophotometer (Thermo Fisher Scientific, Wilmington, DE).

### 2.5 *In vitro* evaluation of *cis*-cleavage activity of Gs12-16 and Gs12-18

The *in vitro cis*-cleavage assay was performed using 1 pmol of Gs12-16, Gs12-18, 2 µL of 10 × rCutsmart buffer, 500 ng of crRNA, and 200 ng of the NDRV S3 gene fragment for each of the different PAMs. The reaction mixture was topped with Diethy pyrocarbonate (DEPC)-treated water to a final volume of 20 µL. The reaction was incubated at 37 °C for 30 minutes. 0.5 µL of Proteinase K (Tiangen, Beijing, China) was added and incubated at 55 °C for 10 minutes to terminate the reaction. The resulting products were then subjected to 2% agarose gel electrophoresis. The difference between the experimental and control groups’ target bands was used to characterize the *in vitro* cleavage activity of different PAM sites. The gray value of the bands was automatically read using Image software, and the cleavage efficiency was calculated according to the following equation: Cleavage Efficiency 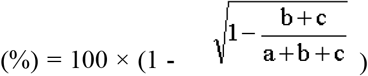, where a represents the wild-type bands, and b and c represent the cleaved bands.

### 2.6 *In vitro* evaluation of *trans*-cleavage activity of Gs12-16 and Gs12-18

The 20 µL reaction mixture for the fluorescence-based assay consisted of 500 nM Gs12-16 or Gs12-18, 1 µM crRNA, 0.75 µM ssDNA-FQ reporter (5’ ROX/GTATCCAGTGCG/3’BHQ2), 2 µL 10 × rCutSmart Buffer and 1 µL double/single-stranded DNA target. The reaction was incubated at 37 °C for 5 minutes and then inactivated at 98 °C for 2 minutes. The solution was exposed to blue light (EP2020, BioTeke) to visualize the reaction products, and images were captured using a smartphone camera. The reaction products were transferred to a 96-well black polystyrene microplate (Costar, Ref: CLS3925) containing 80 µL of nuclease-free water. Fluorescence intensity was measured in a multifunctional microplate reader (EnSpire Multimode Plate Reader, PerkinElmer). The excitation and emission wavelengths for the ssDNA-FQ (5’ ROX/GTATCCAGTGCG/3’BHQ2) reporter were 576 nm and 601 nm, respectively.

### 2.7 Optimization of the effect of temperature and time on the activity of Gs12-16 and Gs12-18

*Cis*-cleavage reactions were performed using PCR-generated DNA of the NDRV S3 gene as the target in a system consisting of 500 nM Gs12-16 or Gs12-18 protein, 1 µM crRNA5, 2 μL 10 × rCutSmart Buffer, 2 μL NDRV S3 dsDNA, and DEPC-treated water up to 20 μL; the negative control omitted crRNA. The *trans*-cleavage reaction was based on this system with the additional 1 µM ssDNA-FQ reporter, and the negative control excluded the dsDNA target. To determine the optimal reaction temperature, reactions were carried out at 25 °C, 37 °C, 45 °C, 55 °C, and 60 °C for 30 minutes. Reactions were performed at 37 °C for 5, 10, 20, 30, and 40 minutes to evaluate the optimal reaction time, respectively. Following *cis*-cleavage reactions, 0.5 μL proteinase K was added to terminate the reaction by incubation at 55 °C for 10 minutes. Subsequent detection was performed using 2% agarose gel electrophoresis. After *trans*-cleavage reactions, inactivation was carried out at 98 °C for 2 minutes, and fluorescence intensity and background noise were measured under blue light using a multifunctional microplate reader.

### 2.8 LAMP amplification of target NDRV S3

Based on LAMP primer design principles, four sets of LAMP primers targeting the NDRV S3 gene sequence were designed using the PrimerExplorer V5 website (Table S2). The amplicon sequences include the best crRNA5-GTTA binding site. The reaction system consisted of 1 μL *Bst* 3.0 DNA polymerase, 2.5 μL 10 × Isothermal buffer, 6 mM MgSO_4_, 14 mM dNTP, a primer mixture (1.6 μM FIP/BIP, 0.2 μM F3/B3, and 0.4 μM LF/LB), 2 μL DNA template, and DEPC-treated water was added up to 25 μL. The templates used were NDRV S3 plasmids with varying copy numbers, while the negative template was DEPC-treated water. The reaction was performed at 65°C for 40 minutes following vortexing, mixing, and instantaneous centrifugation. The amplification products were subsequently identified using a 2% agarose gel.

### 2.9 Sensitivity of CRISPR/Gs12-18 for detection of NDRV S3

The copy number of the pUC19 plasmid containing 662 bp NDRV S3 was calculated based on the molecular weights of the plasmid and the gene as follows: copy number = (plasmid concentration × 6.022 × 10^23^)/(length × 1 × 10^9^ × 650). The constructed pUC19-NDRV S3 plasmid was diluted 10-fold from 3.88 × 10^11^ copies/µL to 3.88 × 10-1 copies/µL for LAMP amplification to evaluate the sensitivity of CRISPR/Gs12-18 for the detection of NDRV S3. The reaction system was as follows: 500 nM Gs12-18, 1 μM crRNA5-GTTA, 2 μL 10 × rCutSmart Buffer, 1 μM ssDNA-FQ reporter, 2 μL LAMP amplification of the target product, 37 °C for 5 min, 98 °C for 2 min, and observe the fluorescence intensity and the background noise in the blue light and Ultraviolet (UV) light.

### 2.10 Statistical Analysis

Statistical analysis was performed using the R programming language. A two-tailed Student’s t-test was used to determine significant differences between treatment and control groups (^***^ *P* < 0.001, ns: not significant).

## 3 Results

### 3.1 *Cis*- and *trans*-cleavage activity of Gs12-16 and Gs12-18 Cas12a orthologs

The sequence similarity between the two novel Cas12a orthologs, Gs12-16 and Gs12-18 nucleases which from our prevous study, was compared with that of other known Cas12a enzymes, such as LbCas12a, FnCas12a, AsCas12a, MbCas12a, and PiCas12a using multiple amino acid sequence alignments. The results showed that the amino acid sequence conservation of Gs12-16 with these five established nucleases was 43.50%, 42.75%, 33.43%, 44.75%, and 39.72%, respectively, while that of Gs12-18 was 47.21%, 43.81%, 34.59%, 45.58%, and 42.14% (Figure 1A).

**Fig. 1.**
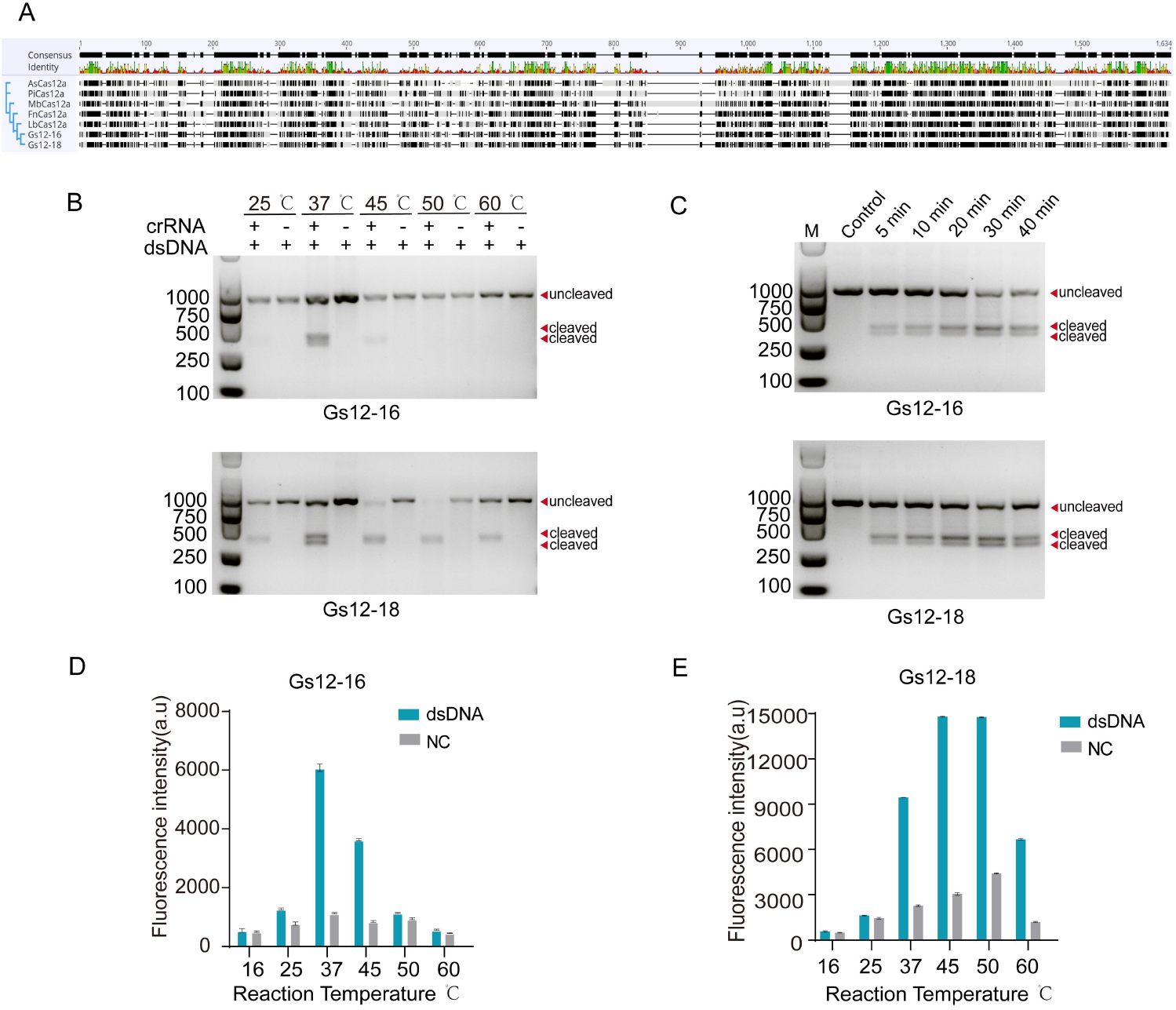
Characterization of Gs12-16 and Gs12-18 nuclease activities. (A) Sequence conservation analysis of Gs12-16 and Gs12-18 against five known Cas12a orthologs. (B) Evaluation of the optimal reaction temperatures for *cis*-cleavage activities of Gs12-16 and Gs12-18 nucleases. (C) Evaluation of the optimal reaction time for *cis*-cleavage activity of Gs12-16 and Gs12-18 nucleases at 37 °C. (D and E) Detection of fluorescence intensity of *trans*-cleavage activities of Gs12-16 and Gs12-18 using a Microplate Reader. dsDNA refers to PCR production of the NDRV S3 fragment, and NC stands for Negative Control.

The Gs12-16 and Gs12-18 proteins were successfully expressed and purified, and the optimal reaction temperatures for *cis*-cleavage activity were evaluated using an NDRV S3 gene fragment as the test target and the PAM sequence as the classical TTTV of LbCas12a separately. The results revealed that Gs12-16 exhibited *cis*-cleavage activity between 37 °C and 45 °C, whereas Gs12-18 demonstrated a more robust temperature tolerance, with activity observed across a broader range of 25 °C to 60 °C, following a 30-minute reaction at five different temperatures (25 °C, 37 °C, 45 °C, 50 °C, and 60 °C) (Figure 1B). We further evaluated the reaction times of Gs12-16 and Gs12-18 nucleases at a fixed temperature of 37 °C. The results showed that both nucleases started to exhibit *cis*-cleavage activity in 5 minutes and reached saturation of *cis*-cleavage activity at 30 minutes (Figure 1C). These results indicated that despite having relatively low sequence similarity with the established Cas12a enzymes, these two novel nucleases exhibited robust *cis*-cleavage activity.

Subsequently, we evaluated the *trans*-cleavage activities of both Gs12-16 and Gs12-18. Our results showed that both nucleases could cleave the ROX-modified ssDNA-FQ reporter. Compared to Gs12-16, Gs12-18 exhibited a relatively wider reaction temperature range of 37–60 °C, with its *trans*-cleavage activity peaking at 37–50°C (Figure 1D and 1E). In contrast, Gs12-16 only displayed *trans*-cleavage activity at 37 °C and 45 °C. Given that Gs12-18 demonstrated *in vitro* cleavage activity across a broad temperature range but exhibited the lowest false positive cleavage activity at 37 °C, we selected 37 °C as the optimal temperature for subsequent experiments (Figure 1E). Both Gs12-16 and Gs12-18 possessed high *cis*- and *trans*-cleavage activities *in vitro*, albeit with different reaction temperatures.

### 3.2 *Cis*-cleavage efficiencies of Gs12-16 and Gs12-18 targeting classical and non-classical PAM sites

To assess and compare the *cis*-cleavage efficiencies of Gs12-16 and Gs12-18, as well as their cleavage preferences for classical (TTV or TTTV) or non-classical PAM sites, we designed 28 crRNAs targeting the NDRV S3 gene fragment, recognizing distinct PAM sequences (Table S1). *In vitro* cleavage reactions were performed using purified Gs12-16 or Gs12-18 proteins, *in vitro* transcribed (IVT) crRNAs, and PCR-amplified NDRV S3 templates. The resulting cleavage products were analyzed by agarose gel electrophoresis (Figure 2A). Our results demonstrate that Gs12-16 and Gs12-18 exhibit efficient *cis*-cleavage activity against approximately 50% of the test target sequences harboring classical PAMs (Figure 2B). Further analysis of the cleavage efficiencies using Image J software revealed that Gs12-18 exhibited relatively higher cleavage activity for most classical PAM-containing sites than Gs12-16 (Figure 2C). Additionally, both Gs12-16 and Gs12-18 could cleave less than 20% of the test targets containing non-classical PAMs (Figure 2D and 2E), with significantly lower cleavage activity observed for these non-classical PAM targets compared to classical PAM-containing targets. These findings suggest that Gs12-16 and Gs12-18 proteins exhibit relatively high *cis*-cleavage activity against classical PAM-containing targets *in vitro*, with Gs12-18 displaying a slightly higher efficiency.

**Figure 2.**
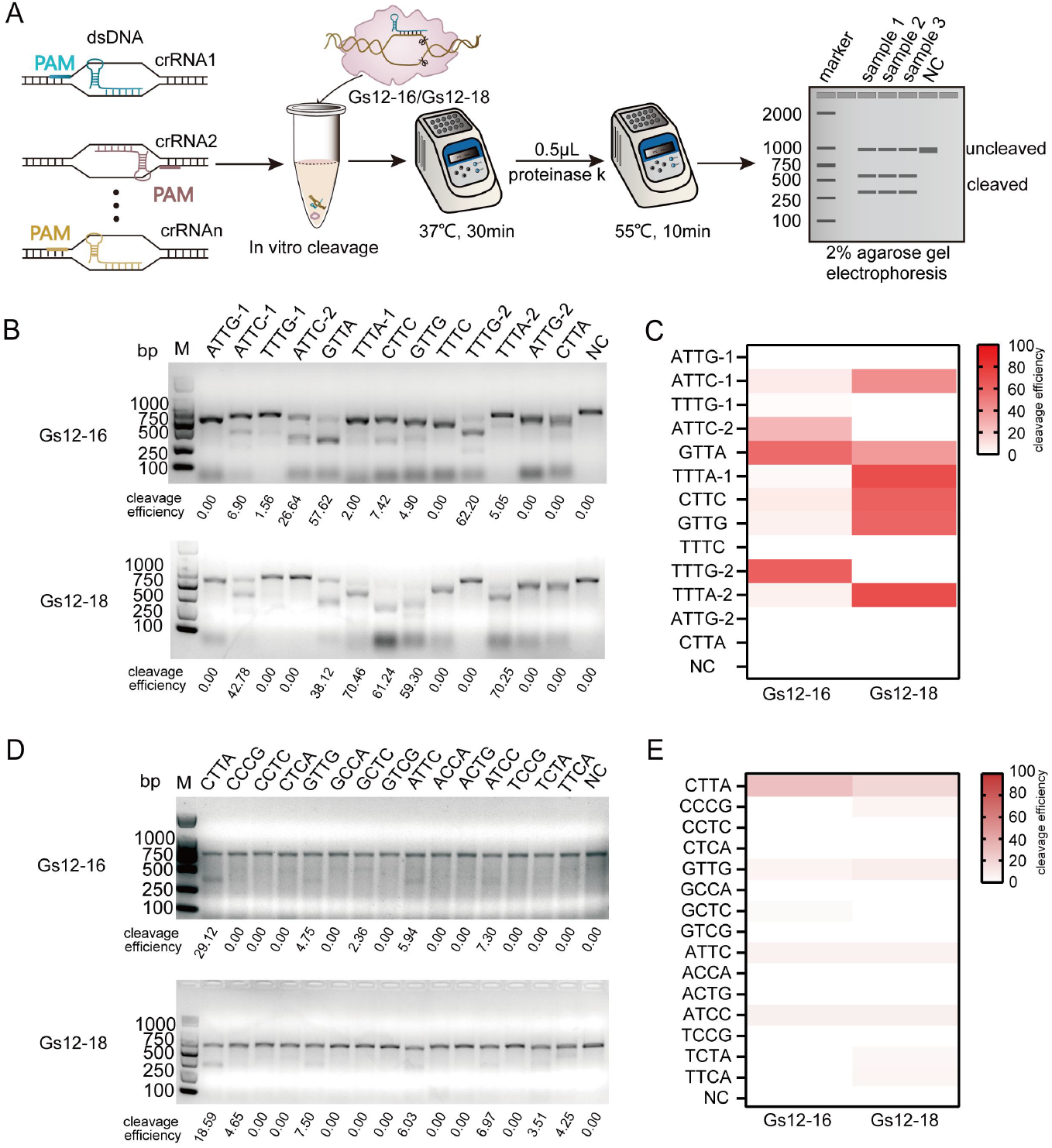
Evaluation of *cis*-cleavage activities of Gs12-16 and Gs12-18 nucleases. (A) Schematic representation of the assay designed to evaluate the *cis*-cleavage activities of Gs12-16 and Gs12-18 nucleases. (B) Agarose gel electrophoresis detection of *cis*-cleavage activity of Gs12-16 and Gs12-18 against classical PAM-containing sites. (C) Heatmap analysis of cleavage efficiencies for classical PAM-containing sites, calculated using Image J software. (D) Agarose gel electrophoresis detection of *cis*-cleavage activity of Gs12-16 and Gs12-18 against non-classical PAM-containing sites. (E) Heatmap analysis of cleavage efficiencies for non-classical PAM-containing sites, calculated using Image J software. Note: NC, negative control; bp, base pairs; M, DNA ladder.

### 3.3 Comparison of *trans*-cleavage efficiencies of Gs12-16 and Gs12-18

To comparatively assess the *trans*-cleavage efficiencies of Gs12-16 and Gs12-18, we employed reactive substrates consisting of dsDNA derived from the amplification product of the NDRV S3 gene fragment, as well as ssDNA test templates generated using matched reverse primers (Figure 3A). A ROX-modified ssDNA-FQ reporter was also utilized to facilitate fluorescent signal detection under blue light using either a blue Light detector or a Microplate Reader (Figure 3A). Our results showed that Gs12-16 and Gs12-18 exhibited similar *trans*-cleavage efficiencies for ssDNA targets harboring classical TTV PAM sites (Figure 3B). However, when evaluating dsDNA targets, the activity of Gs12-18 was relatively higher than that of Gs12-16 (Figure 3C). These findings suggest that Gs12-18 may be a more suitable candidate for CRISPR-based nucleic acid detection applications, owing to their enhanced efficiency in cleaving dsDNA targets.

**Figure 3.**
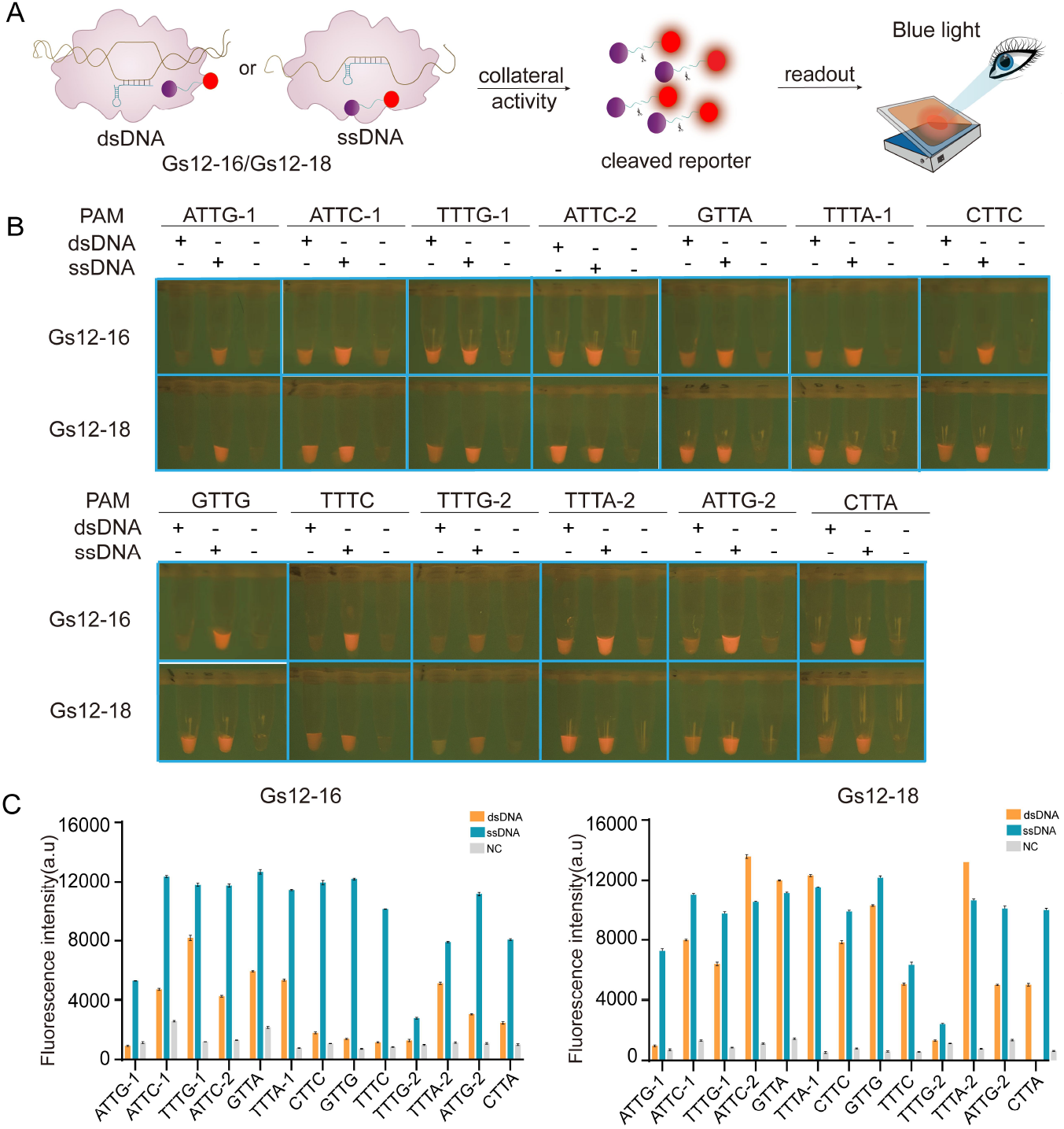
Identification of the *trans*-cleavage activity of Gs12-16 and Gs12-18 nucleases. (A) Schematic diagram illustrating the evaluation of the *trans*-cleavage activity of Gs12-16 and Gs12-18 against classical TTV PAM-containing sites. (B) Visualization of the *trans*-cleavage activities of Gs12-16 and Gs12-18 under blue light. (C) Quantification of fluorescence intensity corresponding to the *trans*-cleavage activities of Gs12-16 and Gs12-18 using a Microplate Reader.

### 3.4 Establishment of a sensitive LAMP-CRISPR/Gs12-18-based nucleic acid detection technology

To achieve highly sensitive, naked-eye visual nucleic acid detection using the Gs12-18 enzyme, we combined it with LAMP. Specifically, we aimed to develop a LAMP-coupled CRISPR-Gs12-18 method for detecting NDRV (Figure 4A). To design a specific crRNA, we used a homology comparison of 26 NDRV S3 gene sequences from the NCBI nr database. The results showed that a crRNA with a PAM of GTTA was conserved among 26 strains (Figure 4B). The activity of this crRNA was also tested (Figure 3B). Next, four sets of LAMP primers were designed (Figure 4C).

**Figure 4.**
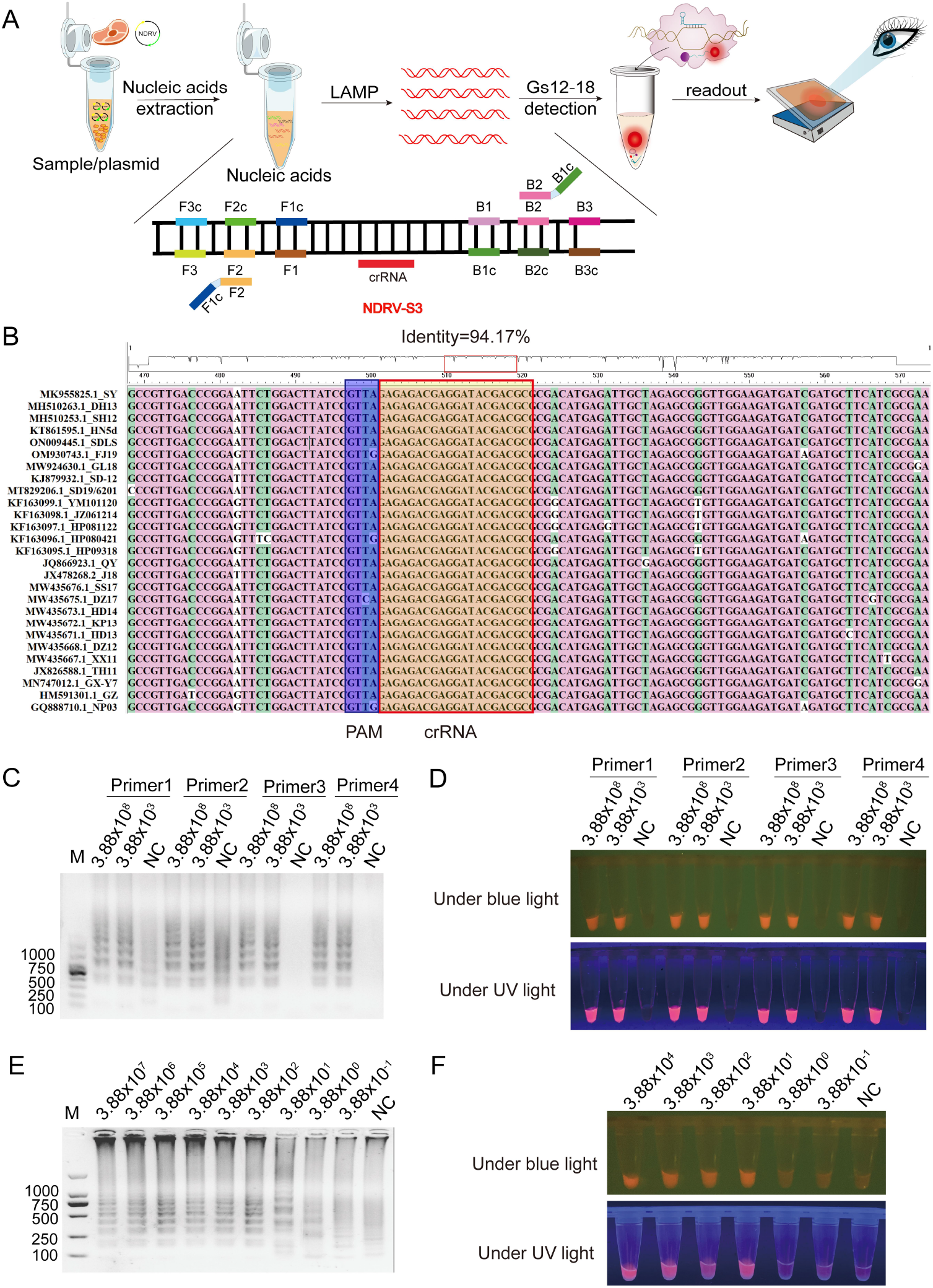
Establishment of LAMP-CRISPR/Gs12-18-based detection of NDRV. (A) Schematic illustration of the LAMP-CRISPR/Gs12-18-based method for visualizing the detection of the NDRV S3 gene. (B) Sequence conservation analysis of candidate highly active crRNA targeting the NDRV S3 gene. PAM, Protospacer Adjacent Motif. (C and D) Identifying optimal LAMP primer pairs for amplifying the NDRV S3 gene fragment and detecting LAMP production using CRISPR/Gs12-18 assays under blue or UV light. (E and F) Evaluation of the limit of detection (LoD) of the NDRV S3 fragment using LAMP or LAMP-CRISPR/Gs12-18 technology.

The template plasmid of NDRV S3 was subjected to 10-fold dilution from 3.88 × 10^8^ to 3.88 × 10^−1^ copies/μL. A LAMP assay for amplification of the NDRV S3 gene was performed, and their products were cleaved using Gs12-18, respectively. The results showed that all four LAMP products could be specifically cleaved by Gs12-18 (Figure 4D). Subsequently, Primer 3 of LAMP was randomly selected to perform follow-up assays. Finally, the limit of detection (LoD) of the LAMP-coupled CRISPR-Gs12-18 method for detecting NDRV S3 was 3.88 × 10^1^ copies/µL (Figure 4E and 4F). These results indicated a highly sensitive and visualized method for detecting NDRV based on LAMP-CRISPR/Gs12-18 technology can be achieved.

## 4 Discussion

The application of RAVI-CRISPR technology for virus detection has emerged as a significant area of research, driven by the need for rapid, accurate, and sensitive diagnostic tools against viral outbreaks and pandemics^[22,24,25]^. This study assessed and compared the *cis*- and *trans*-cleavage activities of two novel Cas12a orthologs, Gs12-16 and Gs12-18, and established a sensitive nucleic acid detection method for NDRV utilizing these enzymes. Our results demonstrated that despite relatively low sequence similarity to known Cas12a enzymes, both Gs12-16 and Gs12-18 exhibited robust *cis*- and *trans*-cleavage activities *in vitro*, with distinct temperature preferences. Gs12-18 showed broader temperature tolerance and higher cleavage efficiency, particularly for dsDNA targets. Furthermore, we successfully developed a highly sensitive and visualized LAMP-CRISPR/Gs12-18-based nucleic acid detection method for NDRV, achieving a LoD of 3.88 × 10^1^ copies/µL.

When comparing the sequence similarity and cleavage activities of Gs12-16 and Gs12-18 to other known Cas12a enzymes, these novel orthologs possess unique characteristics that make them promising candidates for CRISPR-based detection applications^[26]^. The relatively low sequence similarity with known Cas12a enzymes indicates that Gs12-16 and Gs12-18 can provide novel functionalities and applications in gene editing and nucleic acid detection^[23,27,28]^. Their robust *cis*- and *trans*-cleavage activities and distinct temperature preferences offer flexibility in experimental design and application scenarios^[29,30]^. Gs12-18 demonstrated superior performance regarding temperature tolerance and cleavage efficiency, especially for dsDNA targets, which is advantageous for developing reliable and sensitive detection methods^[31]^.

In conclusion, this study effectively assessed and compared the *cis*- and *trans*-cleavage activities of two novel Cas12a orthologs, Gs12-16 and Gs12-18, and established a sensitive LAMP-CRISPR/Gs12-18-based nucleic acid detection method for NDRV. The findings highlight the potential of these novel Cas12a orthologs in broadening the CRISPR toolkit for diverse applications in nucleic acid detection and gene editing. The developed detection method provides a practical solution for the early prevention and control of NDRV, benefiting the global duck farming industry^[32]^.

However, this study also has certain limitations. The experimental data obtained so far may not comprehensively reflect the actual performance of these enzymes in complex biological samples. Further clinical validation is crucial to accurately assess their efficacy and reliability in real-world applications. Additionally, while the LAMP-CRISPR/Gs12-18 method demonstrated high sensitivity, its implementation in field settings requires further optimization and validation to ensure reliability and practicality. Future research will tackle these limitations and explore the full potential of Gs12-16 and Gs12-18 in various CRISPR-based applications.

## CRediT authorship contribution statement

**Yuan Wang**: Writing - review & editing draft, Methodology. **Lanting Fu**: Writing - review original draft, Methodology. **Sheng Li**: Writing – original draft, Software, Methodology. **Dagang Tao**: Writing – original draft, Methodology. **Ping Gong**: Data curation. **Yu Yang**: Data curation. **Shengsong Xie**: Funding acquisition, Writing – review & editing, Supervision. **Cui Wang**: Funding acquisition, review & editing, Supervision. **Daqian He:** Funding acquisition, review, Supervision.

## Acknowledgment

The research was funded by the Donghu High-Tech Zone “open bidding system” project (2023KJB228) cooprated with Wuhan Sunma Biotechnology Corp, Wuhan, Hubei, and the National Natural Science Foundation of China (32302729).

